# Intravital imaging reveals glucose-dependent cilia movement in pancreatic islets *in vivo*

**DOI:** 10.1101/2024.07.03.601879

**Authors:** Olha Melnyk, Jeff Kaihao Guo, Zipeng Alex Li, Jing W. Hughes, Amelia K. Linnemann

## Abstract

Pancreatic islet cells harbor primary cilia, small sensory organelles that detect environmental changes to regulate hormone secretion and intercellular communication. While the sensory and signaling capacity of primary cilia are well-appreciated, it is less recognized that these organelles also possess active motility, including in dense multicellular tissues such as the pancreatic islet. In this manuscript, we use transgenic cilia reporter mice and an intravital imaging approach to quantitate primary cilia dynamics as it occurs in live mouse pancreatic islets. We validate this imaging workflow as suitable for studying islet cilia motion in real time *in vivo* and demonstrate that glucose stimulation corresponds to a change in cilia motility, which may be a physiologic measure of nutrient-dependent fluxes in islet cell function.

## INTRODUCTION

Primary cilia are solitary antenna-like organelles present on the surface of most eukaryotic cells to serve essential sensory and signaling functions^1^. Gene perturbations in ciliary structure, motility, and signaling associate with cellular dysfunction and disease, including metabolic disorders such as obesity and diabetes^1-4^. In the pancreatic islet, functional studies from both human and mouse indicate that primary cilia on islet beta cells receive and integrate a variety of paracrine and juxtacrine signals^5-7^ to modulate glucose-stimulated secretion of insulin, a major hormone required for maintaining systemic energy homeostasis. However, we currently lack experimental evidence as to whether beta cell primary cilia can sense and respond to glucose, as well as the physical characterization of glucose-dependent cilia dynamics, as both the tools and conceptual impetus for investigating such behavior *in vivo* have not been available. Yet, testing this in as a physiologic system as possible is important for understanding the role of primary cilia in regulating normal beta cell function.

Typically defined as non-motile, primary cilia have been classified as strictly sensory organelles whose “9+0” microtubule structure is considered incompatible with axonemal motility^8,9^. This classification sets primary cilia apart from another type of cilia called motile cilia, exemplified by airway epithelial cells and sperm flagella whose coordinated movements constitute their central function. However, primary cilia exhibit considerable diversity in their axonemal structure, as multiple ultrastructural studies including recent 3D volume electron microscopy datasets have demonstrated non-”9+0” microtubule patterns in the primary cilia of pancreatic islets and other tissues^10-16^. Specific to cilia motility, human and mouse islet primary cilia have been found to contain axonemal dynein motors and central pair-associated proteins^17,18^, structural elements that may support active, energy-driven ciliary movement.

Spontaneous primary cilia motion has previously been observed in multiple cell types and postulated as a response to extracellular signals or extraciliary forces^16,19-22^. Functionally, the movement of primary cilia has been observed to accompany changes in intraciliary calcium and cyclic adenosine monophosphate (cAMP) in select experimental systems^23-25^, suggesting a potential link to cellular signaling. We had previously studied beta cell cilia dynamics in intact isolated pancreatic islets *ex vivo*^17,26^ and have demonstrated ciliary motion to be active and regulated by extracellular nutrient level, intracellular ATP, and axonemal dynein^17,18^. Islet cilia movements are irregular and slow compared to those of classic motile cilia but are responsive to ambient glucose and required for beta cell insulin secretion^17^. Meanwhile, the ciliary waveform of beta cell primary cilia has not been quantitatively characterized *in vivo*, thus the question remained whether their observed motility on isolated islets represented true physiologic behavior. It also remains untested whether cilia motility is regulated by glucose *in vivo*, as a measure of cellular response to nutrient flux in the islet environment.

To examine islet cilia dynamics and their glucose responsivity *in vivo*, we used our previously generated Ins1Cre-SSTR3^GFP^ reporter mice^17^ coupled with an intravital microscopy technique developed and optimized for islet imaging in live mouse pancreas^27^. Results are beta cell specific, as expression of the green cilia fluorescent reporter was driven by the Ins1Cre promoter. Retroorbital injection of far red fluorescently conjugated albumin during imaging provided counter-labeling of the pancreatic vasculature, which served as fiduciary landmarks for cilia positional analysis. Intraperitoneal injection of glucose or saline control served as physiologic perturbation. We recorded cilia movements in externalized pancreas of live anesthetized animals and compared changes from baseline to glucose/saline treatments, with each animal serving as its own control before and after treatment. We demonstrate this to be a useful approach for studying islet cilia dynamics and show that beta cell motility can be quantitated *in vivo* and is responsive to physiologically relevant nutrient stimulation in live mice.

## RESULTS

### Live islet cilia imaging using intravital microscopy

The Ins1Cre-SSTR3^GFP^ mouse provided a robust model system for *in vivo* beta cell cilia imaging. Islets within live animal pancreata exhibited robust and stable GFP fluorescence specifically in the ciliary projections of beta cells, visible throughout the depth of the islet, while the vasculature marked by Alexa647-albumin served as a fiduciary for both islet cell and cilia position (**Figure 1**). We chose islets near the pancreas surface for best imaging access and extended the imaging depth using multiphoton microscopy, capturing 1-2 whole islets per pancreas. Within each islet, we visually assessed cilia abundance, distribution, and morphology, which were similar to those previously demonstrated *in vitro* using isolated Ins1Cre-SSTR3^GFP^ islets^17^. Within individual cilia, SSTR3^GFP^ fluorescence expression is stably and uniformly distributed throughout the axoneme, which facilitated waveform tracing and downstream quantitative waveform analysis.

**Figure 1.**
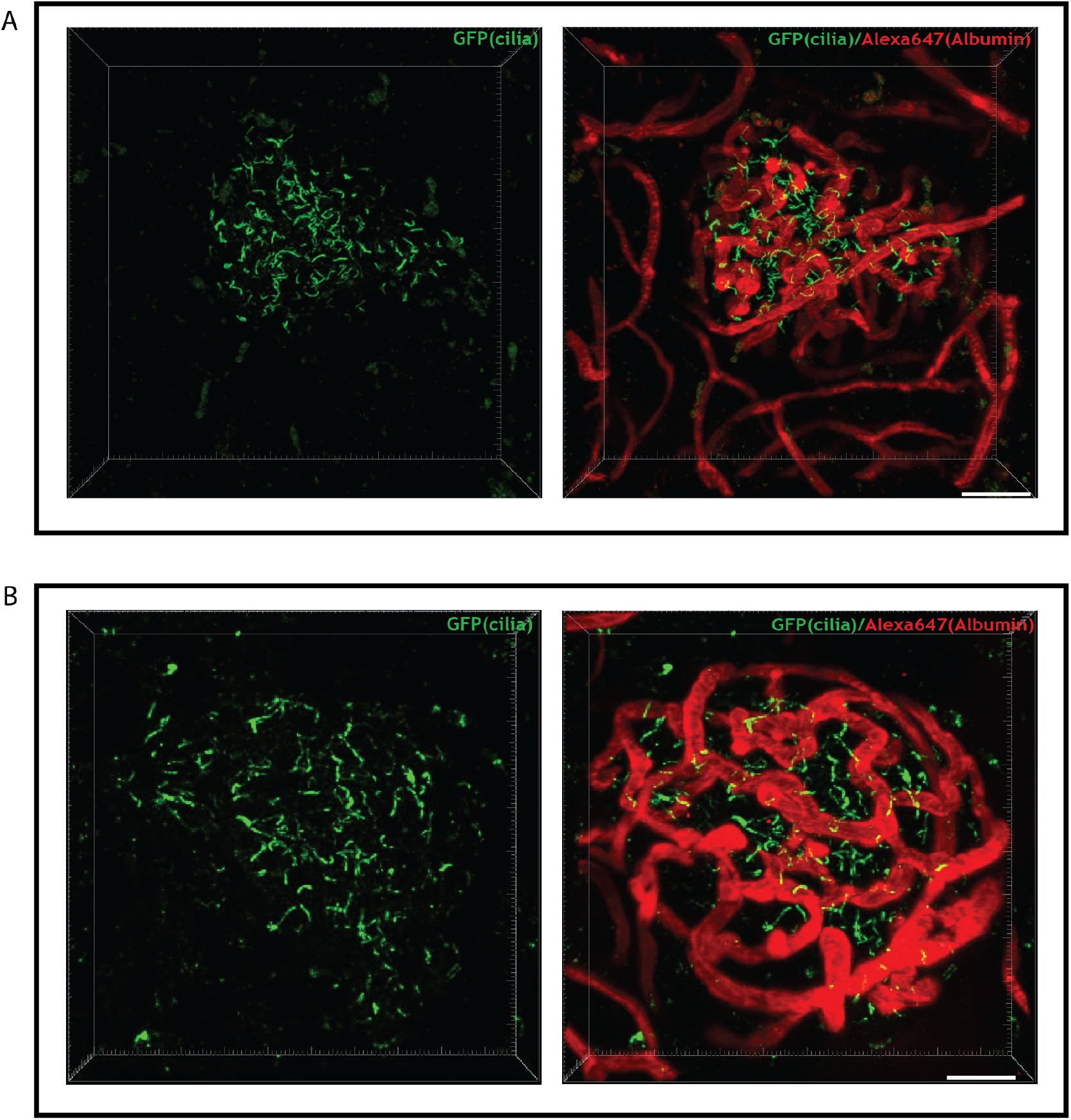
3D visualization of the intravital imaging results from A) Saline injected mouse; B) Glucose injected mouse. Cilia is depicted in green and vasculature network is shown in red. Scale bar = 40 μm.

Imaging parameters for capturing cilia dynamics were optimized both spatially and temporally, based on settings we previously established for imaging cilia *ex vivo*^*17,26*^ and for imaging islet whole-cell dynamics *in vivo*^27^. Specifically, we determined the dimensions of mouse beta cell cilia *in vivo* to be approximately 200 nm in diameter and 2-10 μm in length, similar to prior observations in isolated islets^15,17,26^. Thus, z-stack collection at intervals of 1.5 μm with a total size of approximately 100 μm allowed us to capture all visible fluorescent cilia within the islets. Throughout the imaging duration, we maintained optimal imaging speed and spatial resolution to ensure accurate recognition and capture of cilia (described in more detail in Methods section). The stability of the fluorescent signal further facilitated continuous imaging of cilia motion, both in baseline conditions for 10 minutes and following treatment administration (saline or glucose) for 30 minutes. Imaging settings were kept consistent between experimental conditions to ensure valid comparison of cilia motility changes induced by treatments.

### Beta cell primary cilia exhibit spontaneous motility *in vivo*

We conducted quantitative analysis of cilia waveforms recorded in Ins1Cre-SSTR3^GFP^ mouse islets at baseline and after exposure to saline or glucose bolus (**Supplemental Videos S1-S4, Supplemental Figure 1**), and as shown in **Figure 2 (A)**. Individual cilia were identified and isolated in the images (**Figure 2 (B), (D)**), and their positions were evaluated over time in response to saline (**Figure 2 (C)**) or glucose (**Figure 2 (E)**) injection. Quantification of these movements revealed that most cilia exhibited slow, irregular, small-amplitude fluctuations at the baseline physiologic state (**Figure 3**). Cilia movement was observed throughout the islet including the islet interior which is densely cellular and considered a constrained physical space, but nonetheless permitted cilia movement within the interstitium. Waveform analysis performed here showed that the average values for motion characteristics at baseline were comparable to those observed in *ex vivo* studies^17^. Across different animals, the general motion of cilia appeared consistent, with variations in frequency and amplitude suggesting a lack of discernible patterns or periodicity in baseline motion. Specifically, *in vivo*, we observed a range of cilia wave frequencies across different islets and animals (mean ± SEM = 0.01050 ± 0.0005609 Hz, N = 7 mice/islets, n = 70 cilia) and amplitude (mean ± SEM = 0.3011 ± 0.02104 rad), suggesting that mouse beta cell cilia exhibit heterogeneous motile behavior under normal physiologic conditions.

**Figure 2.**
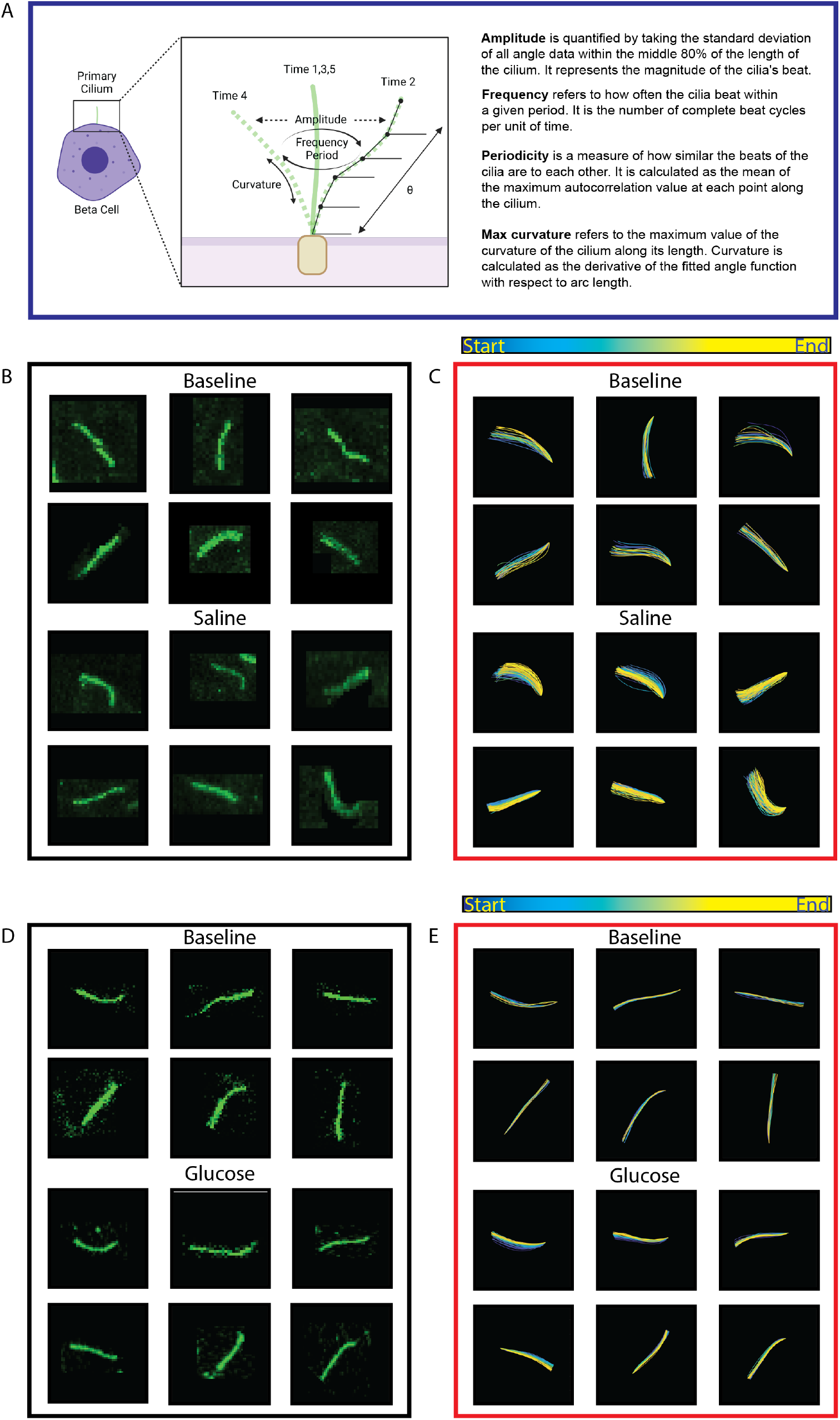
(A) Schematic and definitions of the waveform analysis parameters. (B) Representative individual cilia tracings used for data analysis of saline injected mouse. (C) Waveform tracings depicted in color heatmaps in MatLab for saline injected mouse. (D) Representative individual cilia tracings used for data analysis of glucose injected mouse. (E) Waveform tracings depicted in color heatmaps in MatLab for glucose injected mouse. Total number of mice N = 4 saline, 3 glucose; 10 cilia analyzed per mouse.

**Figure 3.**
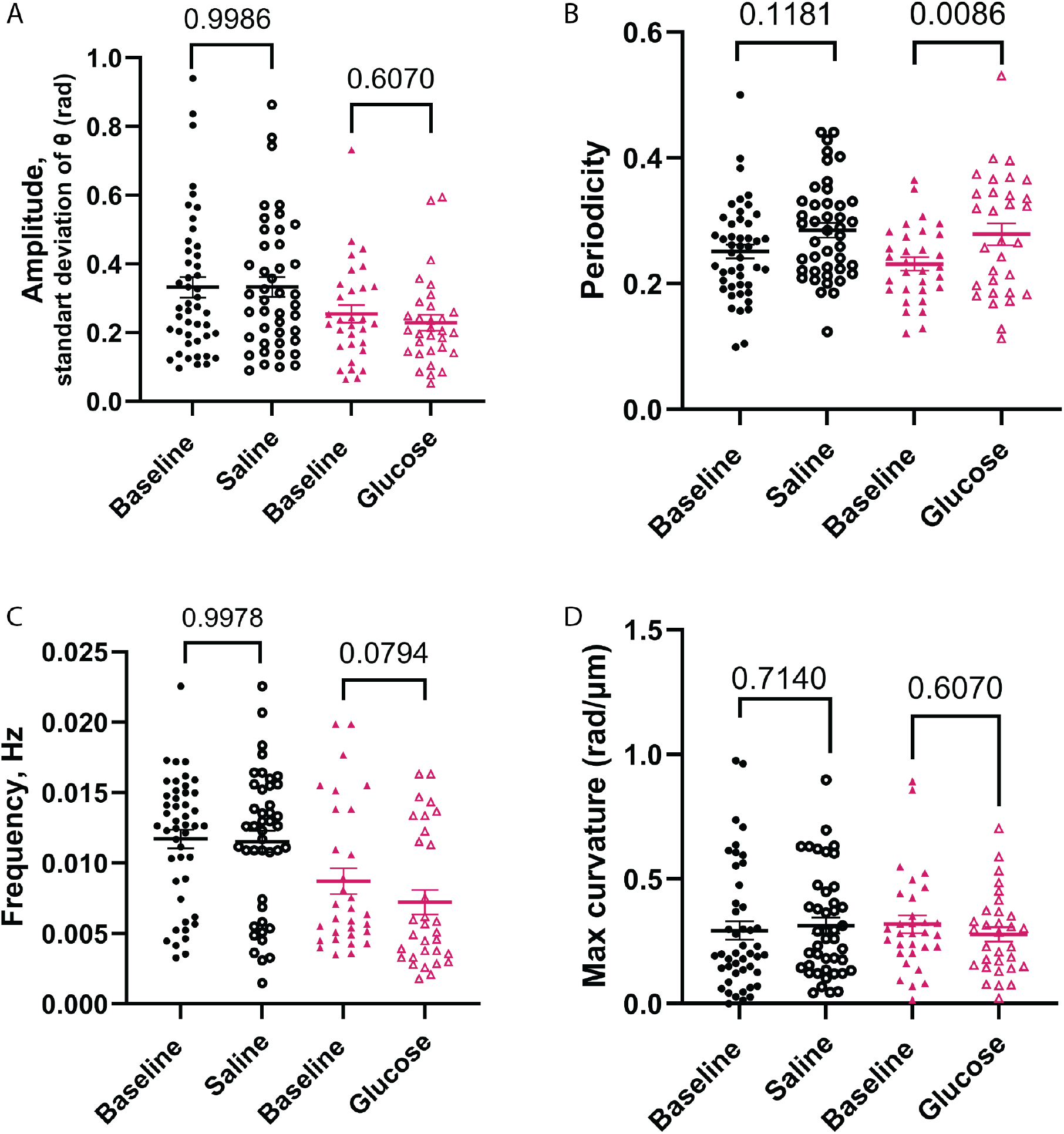
Waveform analysis shows glucose-dependent changes in beta cell cilia motility *in vivo*. Composite results showing saline injected mice (black) and glucose injected mice (pink). Each dot indicates the measured absolute value of individual cilia identified in an islet of imaged mice (either saline or glucose injected). The four waveform parameters analyzed included: (A) amplitude, (B) periodicity, (C) frequency, and (D) maximum curvature. N = 4 (saline) and 3 (glucose) mice, n = 10 cilia/mice, each individual p value is shown on the graphs.

### Beta cell cilia motility changes in response to glucose stimulation

To test whether cilia motion responds to physiologic beta cell stimulation, we injected Ins1Cre-SSTR3^GFP^ mice intraperitoneally with glucose (2 g/kg) or saline vehicle (of equal volume) during intravital imaging and compared cilia waveforms between treatments. A 10-minute segment of imaging was performed pre-injection as baseline to compare with post-injection cilia responses. Each mouse served as its own control pre- and post-injection to control for variability in individual responses to treatment and animal to animal variability in cilia motion. Saline injection (compared to baseline no-injection) did not change any parameters of cilia waveform, including amplitude, periodicity, frequency, and curvature (**Figure 3A, 3; Supplemental Video S1, S3**), ruling out IP injection itself as a confounding factor in altering ciliary motion. In contrast to saline, glucose injection stimulated cilia motion (**Figure 3B, 3; Supplemental Video S2, S4**). While there were no observable changes in amplitude (**Figure 4A**), glucose stimulated a significant increase (*p* = 0.0086) in cilia periodicity (**Figure 4B**), a measure of waveform coordination and regularity, and a near-significant reduction (*p* = 0.0794) in ciliary oscillation frequency (**Figure 4C**). Cilia curvature remained overall unchanged (**Figure 4D**). Analysis for individual mice is shown in **Supplemental Figure 2**. These results indicate that cilia moved with greater coordination and regularity with glucose stimulation. Wave amplitude and cilia curvature also did not change significantly in our analyses, isolating periodicity as the most salient response of mouse beta cell cilia to glucose stimulation.

## DISCUSSION

Primary cilia motility is an observed phenomenon across cell and tissue types^17,19,20^, while quantitative characterization and mechanistic understanding for their motile behavior have been lacking. Prior observations from *in vitro* and *ex vivo* systems may also incompletely capture physiologic cilia dynamics due to artificial experimental conditions. To address these challenges, we used intravital microscopy in a beta cell cilia-GFP reporter mouse model^26^ to study islet cilia movement directly in the live mouse pancreas. Our results show that a) beta cell primary cilia exhibit spontaneous and constant motion under physiologic conditions *in vivo*, and b) cilia motility is responsive to blood glucose changes, as indicated by increased cilia waveform periodicity following glucose injection in healthy live mice.

These results demonstrate a novel aspect of physiologic beta cell glucose responsiveness in the form of primary cilia motility, consistent with the known role of cilia as glucose-dependent regulators of beta cell function.

Our study addresses the question of whether pancreatic islets as a physical environment could support cilia movement and whether cilia do in fact move within intact islets. The answers are yes, and yes. In support of our direct observations by live cilia imaging, volumetric electron microscopy studies in both rodent and human islets^14,16,28^ have demonstrated cilia projection into intercellular spaces which, albeit limited in scale, can support ciliary movement over micron-range distances and interactions with neighboring cilia, as demonstrated by electron microscopy^14,16,28^. Functionally, these intraislet spaces are an important site of nutrient and paracrine flow that provides signals for ciliary sensing and communication. Thus, the glucose-responsive *in vivo* motility characterized in our study likely represents a physiologic property of islet primary cilia that contributes to their sensing and signaling function.

Biomechanically, primary cilia motility differs from that of motile cilia. Our *in vivo* islet cilia waveform analyses, along with prior experimental observations in non-islet systems^20,21,29^, show that primary cilia oscillate in irregular patterns in contrast to the synchronized beating of motile cilia. This difference in waveform regularity may be explainable by the dynamic and variable cytoskeletal structure of primary cilia, particularly their lack of stable central microtubule pairs that are considered required for producing coordinated waveforms^12,13,16^. Nonetheless, subunits of axonemal dynein and other motor proteins have been identified in human and mouse islet primary cilia by light and electron microscopy, along with ultrastructural imaging showing transient functional central pairs and linkers between microtubule doublets^14,18^. Thus, primary cilia may contain structural motifs to support an alternate, if not moderated, capacity for movement in adaptation to their unique environment.

Our data shows that islet cilia movement is intrinsic to their behavior, as has been observed in previous experimental systems, and may serve a functional purpose in islet cell responses to glucose. What our study does not distinguish is whether islet cilia movements are driven by active, cilia-intrinsic forces e.g. axonemal dynein, or by external forces such as shear stress from the extracellular matrix^29-31^, the cortical actin network^32,33^, and fluid flow from the vasculature network within the islets^34,35^, or a combination thereof^20,36^. Without direct experimental data, we speculate that both internal and external forces may be at play, and that the biomechanics and anatomy of the pancreatic islet may collectively influence cilia motion, subject to change from healthy to diseased pancreatic states. Our study also does not address whether beta cell cilia movement *in vivo* is required for insulin secretion, an important functional question that should be addressable in future studies using dynein mutant mice combined with intravital imaging of cilia motility, calcium imaging, and insulin secretion events.

The novelty of this study lies in its combination of cilia reporter animals with intravital imaging technology, which allows for the monitoring of primary cilia dynamics *in situ*. Despite its potential, the intravital imaging approach also has notable caveats. It is low-throughput, sampling only one or two islets per animal and is limited to superficial islets on the pancreas surface, leaving deeper islets inaccessible. Additionally, it is prone to artifacts from animal movement and breathing, necessitating post-imaging stabilization and corrections. Despite these limitations, we find it a suitable method for studying cilia dynamics *in vivo* and one that should open doors for follow-up studies of primary cilia motility across tissue types. The use of an insulin promoter-driven cilia reporter model offers beta cell-specific interrogation, a feature that could be replicated in future studies with virally delivered cilia biosensors. While the study provides a method to observe cilia dynamics *in vivo*, it also opens new research questions about the mechanism and physiologic significance of primary cilia movement, such as how cilia motility might augment islet cell nutrient sensing and communication. Future research should examine the generalizability of primary cilia motility in other cell types and tissues, the structure-function relationships underlying axonemal dynein and cilia waveform generation, and the bioenergetic control of cilia beating. Only then can we truly tease apart the functional role of beta cell primary cilia motility in systemic metabolic response.

## Supporting information

Supplemental Video 1_Saline Baseline

Supplemental Video 2_Glucose Baseline

Supplemental Video 3_Saline

Supplemental Video 4_Glucose

## ACKNOWLEDGMENTS

We gratefully acknowledge the development of the cilia autotrace protocol by Louis Woodhams in the laboratory of Philip Bayly. Parental strain for the SSTR3-GFP reporter mouse was a gift from Bradley Yoder at University of Alabama Birmingham. We thank Jeong Hun Jo from the Hughes laboratory for mouse husbandry, genotyping, and phenotyping. We acknowledge the support of the Indiana Center for Biological Microscopy and the Optical Microscopy Core of the Indiana Diabetes Research Center (NIH P30-DK-097512) for intravital imaging. This work was supported by National Institutes of Health grants DK115795 to JWH and intravital imaging support from R01-DK124380 and R03-DK115990 to AKL.

## AUTHOR CONTRIBUTIONS

Conceptualization, J.W.H. and A.K.L.; Methodology, O.M., Z.A.L., J.W.H., A.K.L.; Investigation, J.K.G., O.M. Z.A.L, and S.Y.W.; Writing – Original Draft, J.K.G.; Writing – Review & Editing, O.M., J.W.H., and A.K.L.

## SUPPLEMENTAL INFORMATION TITLES AND LEGENDS

**Supplemental Figure 1**. 3D reconstruction of the imaged cilia. Cilia which motion was analyzed, depicted in yellow.

**Supplemental Figure 2**. Waveform parameters for individual saline injected mice (gray) and glucose injected mice (red). (A) Each dot indicates the computed frequency of one cilium identified from individual mice found in both the baseline and injection time period. (B) Same as (A) for periodicity. (C) Same as (A) for amplitude (standard deviation of tangent angles to cilia). (D) Same as (A) for wavelength; N = 4 (saline) and 3 (glucose) mice, n = 10 cilia/mice.

**Supplemental Figure 2**. Schematic of the experimental methods used.

Video S1. Baseline imaging of the islet

Video S2. Imagin post saline injection

Video S3. Baseline imaging of the 2^nd^ islet

Video S3. Imaging post glucose bolus.

## METHODS

### Intravital Imaging of Mouse Pancreas

Beta cell-specific fluorescent cilia reporter mice were generated by crossing Ins1Cre (JAX 026801) and SSTR3^GFP^ (JAX 024540) parental strains^22^. Mice (males and females, 58 weeks of age) were anesthetized using inhaled isoflurane (2.5-3%) prior to imaging procedures. The pancreas was externalized through minor surgical intervention as we have previously described ^27^. Briefly, a small incision was made on the left side of the mouse abdomen to carefully expose the pancreas, which was then gently positioned on the surface of the imaging dish. Surrounding the pancreas, a supportive saline solution was administered. Subsequently, the mice were positioned on a heated microscope stage and covered with a heating blanket to maintain body temperature. Continual monitoring of body temperature was conducted via an anal probe to ensure a comfortable environment throughout the procedure. Through the microscope eyepiece, identification of the islet within the pancreas was facilitated by the presence of the green fluorescent protein (GFP) signal emitted from the beta cell cilia. Intravital imaging of pancreatic islets at various depths was conducted using a multi-photon Leica TCS SP8 DIVE microscope equipped with a 25x water objective. The excitation of cilia fluorescence was achieved using a 920 nm Spectra-Physics MaiTai DeepSee laser, with emission collected within the 500nm-550nm spectral window using the HyD hybrid detector. To visualize the vascular network of the islet, mice were administered Alexa Fluor 647-conjugated Albumin. Excitation of Albumin was performed with an 840nm laser, and emission was detected within the 650nm-700nm range using the HyD hybrid detector. In addition, the emission from the same laser was collected within the 500nm-550nm spectral window, to enable noise/background identification from the cilia imaging. Images were collected at a scanner frequency of 600Hz, resulting in 512×512 pixel images collected with a time resolution of 1.88 frames per second. Each mouse initially underwent a 10-minute baseline imaging session, followed by either an IP injection of saline or glucose (2 mg/g), and subsequent imaging for 30 minutes to observe longitudinal cilia motion and responses. Imaging and subsequent waveform analysis approach is summarized in **Supplemental Figure 2**.

### Image Processing and Analysis

Following data acquisition, the raw data were imported into FIJI^37^ for image analysis and converted into 8-bit files. For each video containing an imaged islet either pre- or post-IP injection, the channel containing the background fluorescence signal was subtracted from the GFP channel to obtain a background-corrected channel with the true fluorescence signal along with the red vasculature channel, if available. This combined file was imported into Imaris Viewer 10.0.1 for 3D reconstruction and visualization of the islet. Using the 3D reconstruction, cilia were identified and selected for further analysis based on the following criteria:

- The cilium is fully found in the imaging frame of view with sufficient fluorescence signal for the full duration of both the baseline and injection time periods;
- The cilium can be easily isolated and does not intersect substantially with other cilia;
- The cilium does not exhibit excessive curvature;

Each selected cilium identified in Imaris Viewer was matched with its same position in the 4D FIJI file based on its shape and location within the islet and recorded for reference. In FIJI, for each cilium, the range of cross-sections where the given cilium could be identified at any time point were recorded. A maximum intensity projection was performed using only the given range of cross-sections, forming a time lapse video of each cilium, which was subsequently stabilized using the Correct 3D Drift plugin available in FIJI. Additional cropping was performed frame by frame to remove fluorescence signals not belonging to the cilium of interest. This file was then saved and passed into a MATLAB script^17^, which identifies the cilium’s position over time and calculates waveform parameters describing its trajectory. For each cilium’s video, the position of the cilium was manually traced from base to tip, where the base was identified as the end of the cilium that remained more stationary and exhibited less dynamic remodeling over time (Figure 1). The validity of the cilium’s motion trajectory was assessed, and the position, angle, frequency, periodicity, amplitude, and wavelength data were recorded for all cilia. This process was then repeated for all identified cilia from islets (n = 10 cilia per islet) in both baseline and injection data collection periods for saline (N = 4) and glucose (N = 3) injected mice.

### Statistical Analysis

Computed data points for frequency, periodicity, amplitude, and wavelength were pooled across all mice from pre- and post-IP injection experimental periods and normalized to the mean value of the pre-IP injection time period that had the same experimental treatment (either saline or glucose) to obtain fold changes over the median baseline values. A Student’s t-test was performed to compare the fold changes between the baseline and experimental conditions for both the saline and glucose injected mice. The same procedure was performed using data from individual mice to obtain fold changes and a t-test was used (**Figure 3**).

---

**Supplemental Figure 1.**
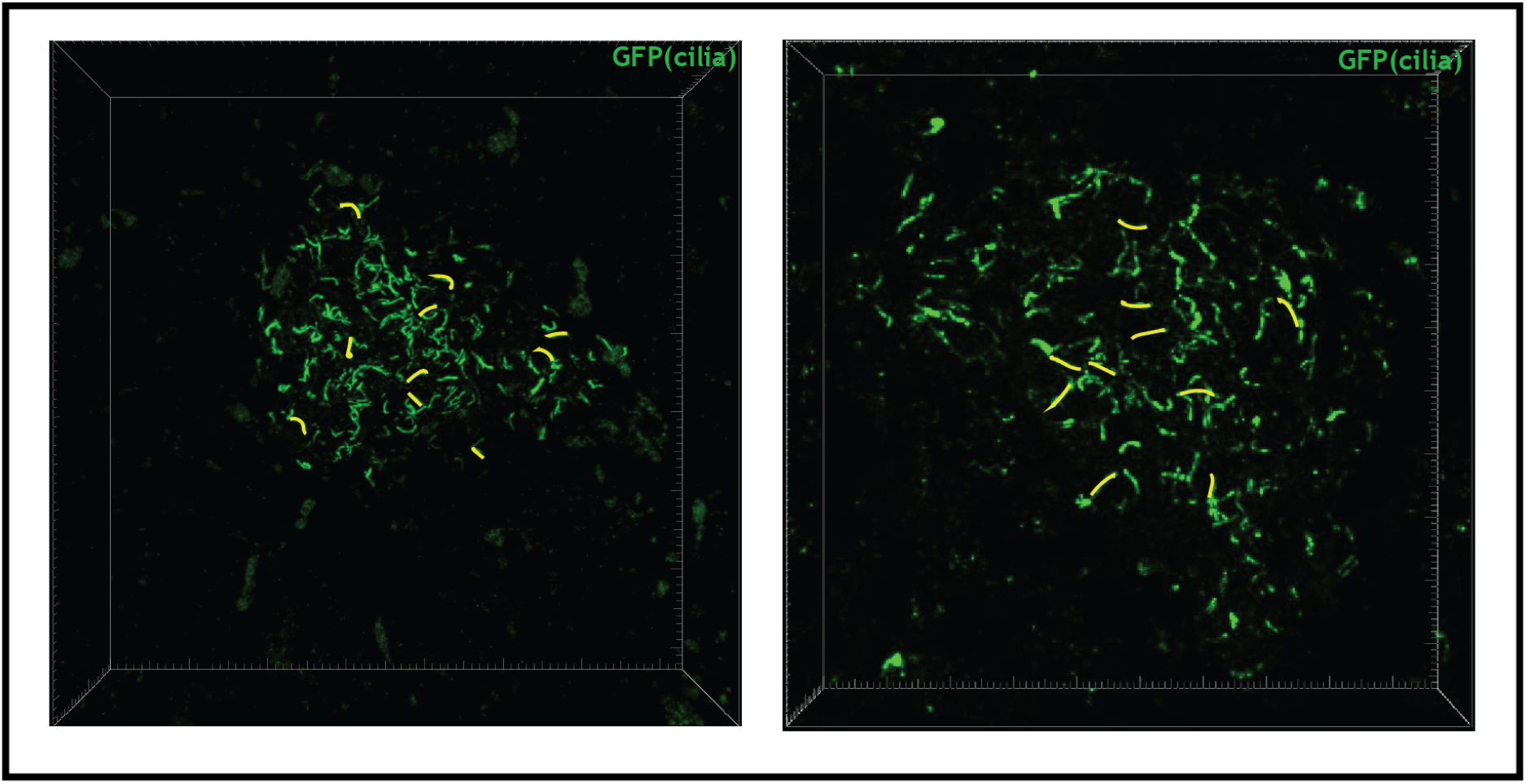
3D reconstruction of the imaged cilia. Cilia which motion was analyzed, depicted in yellow.

**Supplemental Figure 2.**
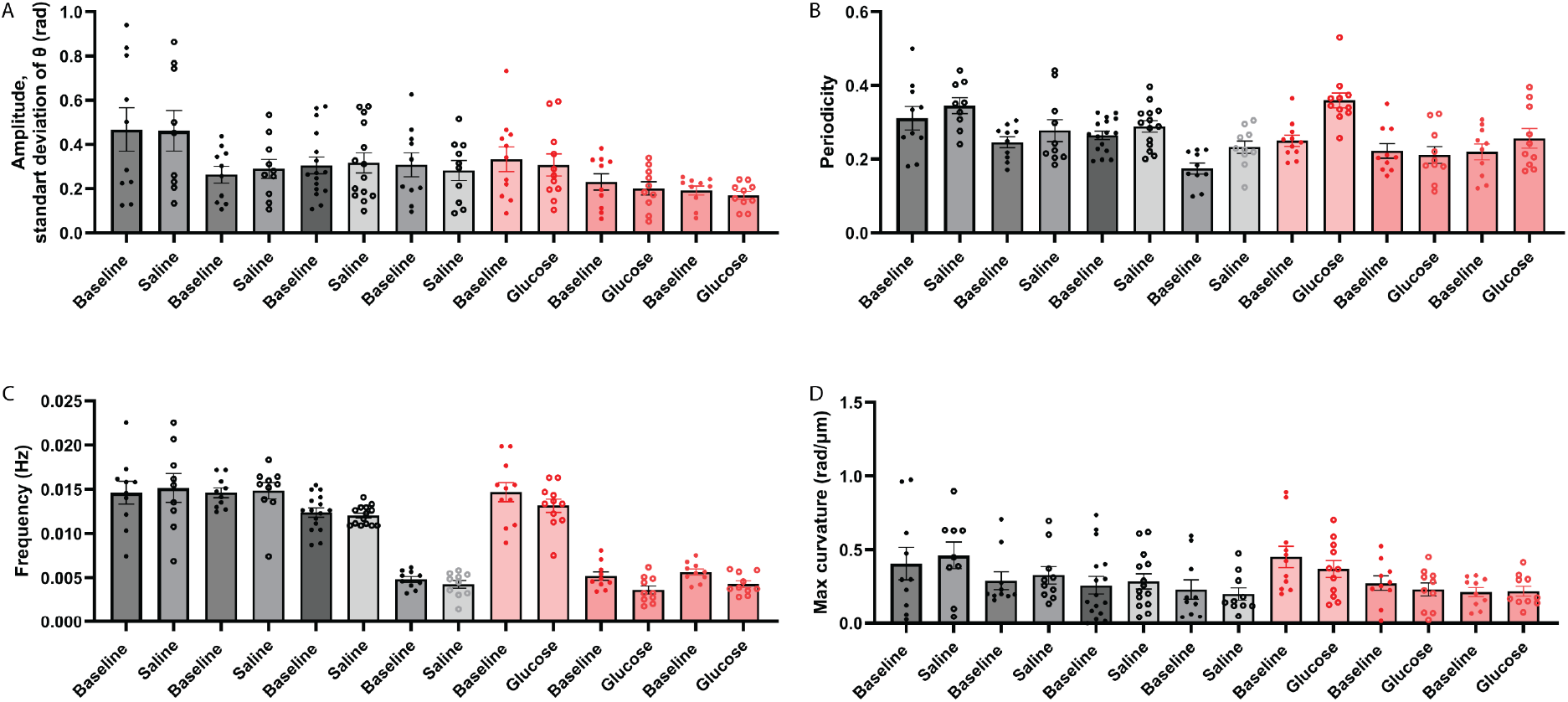
Waveform parameters for individual saline injected mice (gray) and glucose injected mice (red). (A) Each dot indicates the computed frequency of one cilium identified from individual mice found in both the baseline and injection time period. (B) Same as (A) for periodicity. (C) Same as (A) for amplitude (standard deviation of tangent angles to cilia). (D) Same as (A) for wavelength; N = 4 (saline) and 3 (glucose) mice, n = 10 cilia/mice.

**Supplemental Figure 3.**
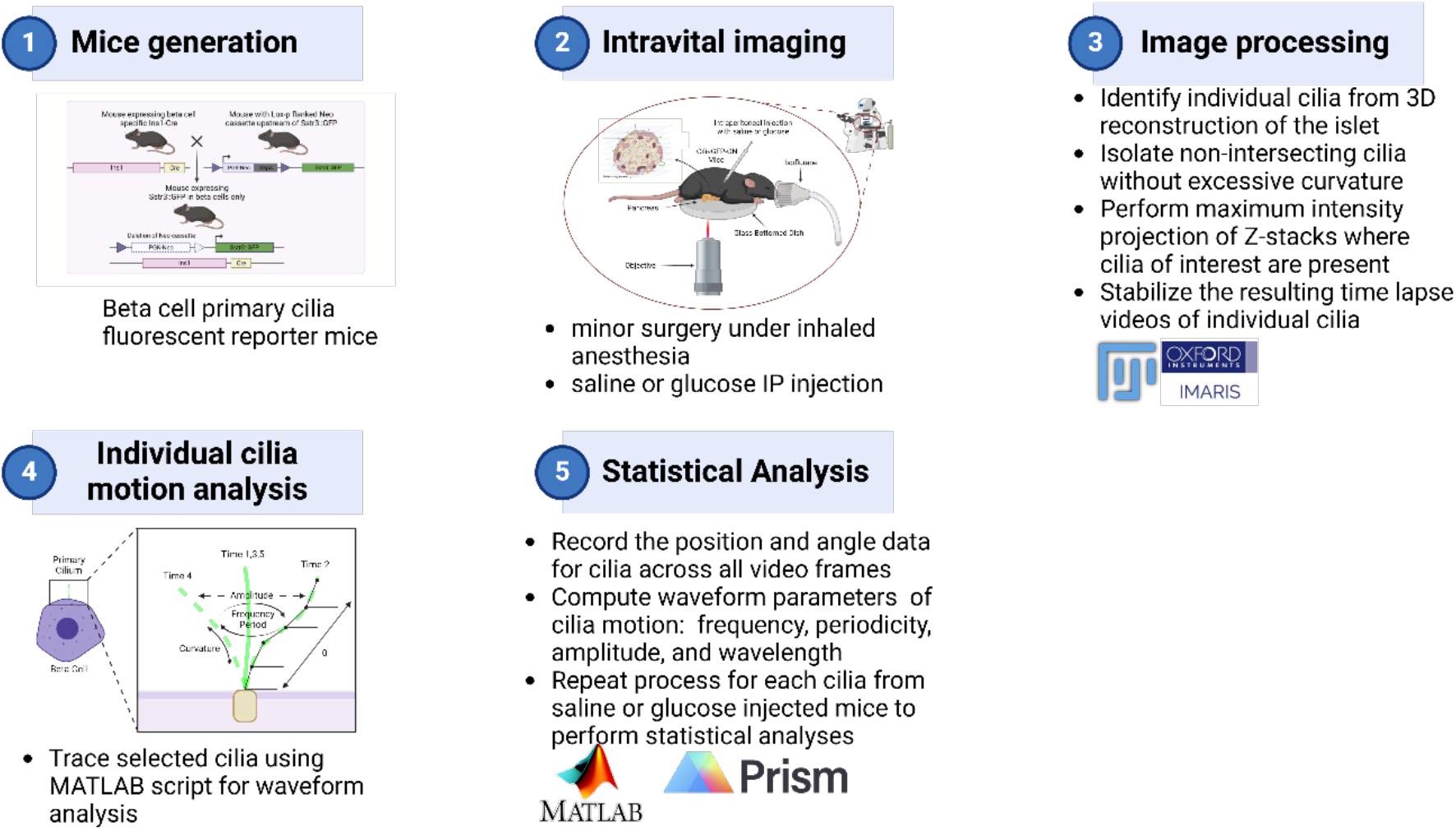
Schematic of the experimental methods used.

